# Storage cell proliferation during somatic growth establishes that tardigrades are not eutelic organisms

**DOI:** 10.1101/2023.10.12.562009

**Authors:** Gonzalo Quiroga-Artigas, María Moriel-Carretero

## Abstract

Tardigrades, microscopic ecdysozoans renowned for their resilience to extreme environments, have long been thought to maintain a constant cell number after completing embryonic development, a phenomenon known as eutely. However, sporadic reports of dividing cells have raised questions about this assumption. In this study, we investigated whether tardigrades truly exhibit a fixed cell number during somatic growth using the model species *Hypsibius exemplaris*. Comparing hatchlings to adults, we observed an overall increase in the number of storage cells, a tardigrade cell type involved in nutrient storage. To assess cell proliferation, we monitored DNA replication via the incorporation of the thymidine analog 5-ethynyl-2’-deoxyuridine (EdU). A significantly higher number of storage cells incorporated EdU while animals were still growing. Starvation halted both animal growth and storage cell proliferation, linking the two processes. Additionally, we found that EdU incorporation in storage cells is associated with molting, a critical process in tardigrade post-embryonic development, since it involves cuticle renewal to enable further growth. Finally, we show that hydroxyurea, a drug that slows down DNA replication progression, strongly reduces the number of EdU^+^ cells and results in molting-related fatalities. Our data not only provide a comprehensive picture of replication events during tardigrade growth but also highlight the critical role of proper DNA replication in tardigrade molting and survival. This study definitively challenges the notion of eutely in tardigrades, offering promising avenues for exploring cell cycle, replication stress, and DNA damage management in these remarkable creatures as genetic manipulation techniques emerge within the tardigrade research field.

**SIGNIFICANCE:** Tardigrades, microscopic invertebrate animals renowned for their resilience in extreme conditions, have traditionally been considered eutelic, implying little to no somatic cell proliferation during their growth. However, a few isolated reports challenged this notion. In this study, using the emerging model *Hypsibius exemplaris*, we provide unequivocal molecular evidence of DNA replication and proliferation in a specific tardigrade cell type called ‘storage cells’, primarily involved in nutrient storage, throughout the animal’s growth. Furthermore, we demonstrate that this proliferation is associated with the timing of cuticle molting, and we highlight the critical role of proper DNA replication in tardigrade molting and survival. Our research definitively resolves the long-standing controversy surrounding tardigrade eutely, opening up uncharted territories in tardigrade research.

## INTRODUCTION

Tardigrades, often called water bears, are tiny animals (0.1-1mm long) that belong to the superphylum Ecdysozoa (Fig. 1A). This clade is characterized by the presence of an exoskeleton or a cuticle, which acts as a protection for their bodies (1). Tardigrades’ cuticle plays pivotal roles in their life cycle, enabling them to respond and adapt to diverse environmental challenges (2). Like other ecdysozoans, tardigrades must undergo a process called ecdysis (*i.e.,* molting), which involves producing a new cuticle and shedding the old one (known as exuvium), in order to grow in size (1, 2). Tardigrades are renowned for their exceptional capacity to withstand extreme conditions, leading to a significant surge in research on this subject in recent years (3, 4). However, a more comprehensive understanding of tardigrade biology under physiological conditions is also crucial, as it is necessary to really understand the changes they undergo when exposed to harsh scenarios.

**Figure 1.**
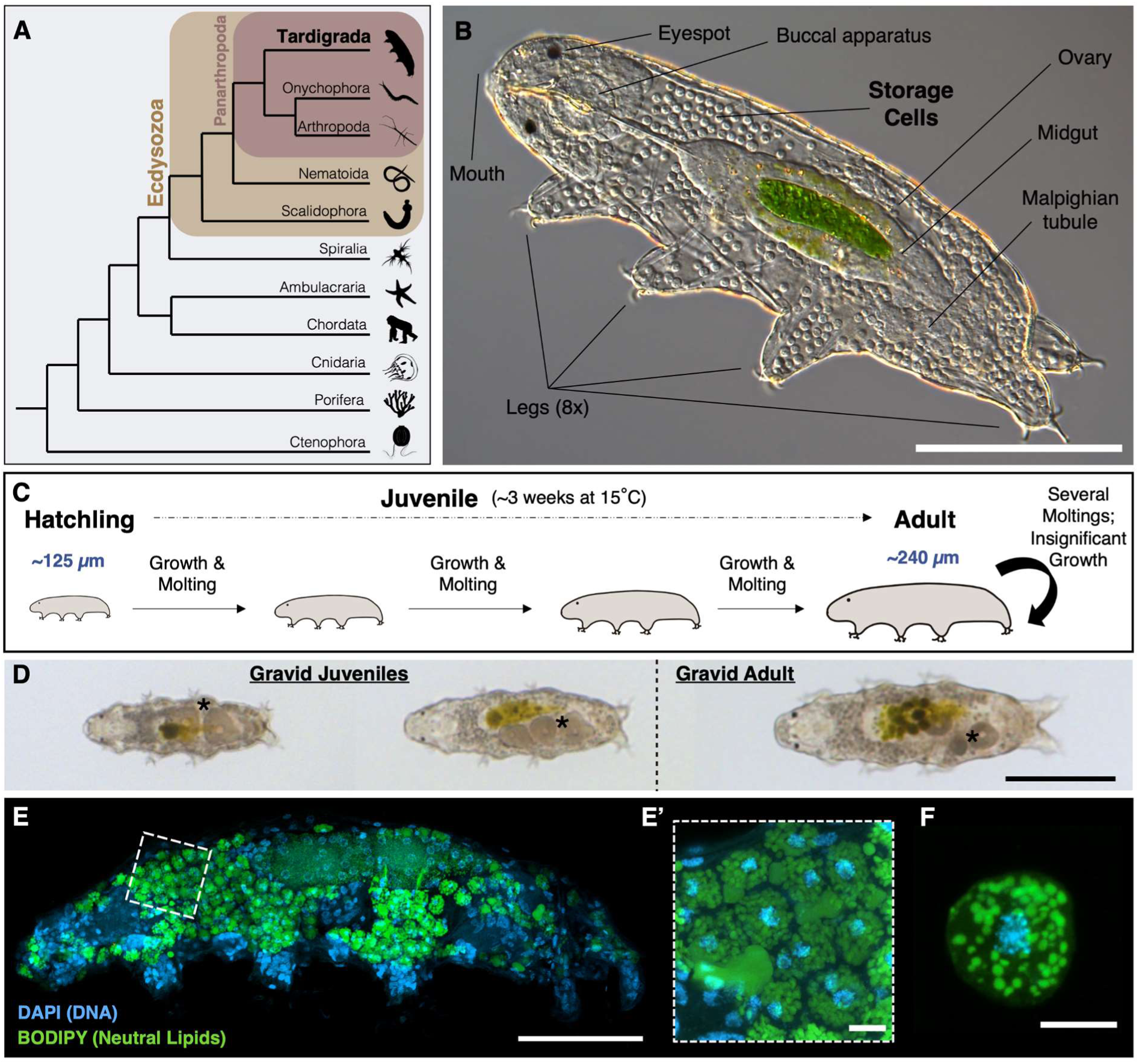
The tardigrade *Hypsibius exemplaris* and its storage cells. (A) A simplified animal phylogeny, highlighting the position of tardigrades (Tardigrada) within the Ecdysozoa. Tardigrades share common ancestry with the Onychophora + Arthropoda group, collectively forming the Panarthropoda clade (after Schultz *et al*. (51), Wu *et al.* (52), Laumer *et al*. (53), Dunn *et al*. (54), and Campbell *et al*. (55)). Certain groups like Placozoa and Xenacoelomorpha are omitted for simplicity. Images sourced from Phylopic.org. (B) Anatomy of *H. exemplaris*. Internal organs are revealed in a smushed specimen. The cuticle refracts light and envelops the animal, while the green patch inside the midgut indicates the digestion of *Chlorococcum* algae. (C) Schematic representation of *H. exemplaris* post-embryonic development. (D) Images of two gravid juveniles at different sizes and a gravid fully-grown adult. Black asterisks indicate the location of oocytes. (E) Representative image of a gravid juvenile showing cell nuclei (DAPI, blue) and neutral lipids (BODIPY, green), primarily visible in lipid-rich storage cells. (E’) Magnification of the region outlined in (E), emphasizing lipid-filled storage cells. (F) One storage cell isolated from the animal, stained as in (E). The full depth of the tardigrade (E) and of the storage cell (F) was captured in confocal z-stacks, and the images shown are maximum projections. In (E’), a sub-stack was used to facilitate lipid droplet visualization in the storage cells’ cytoplasm. Scale bars: 100 µm in (B, D); 50 µm in (E); 5 µm in (E’-F).

Among tardigrades, *Hypsibius exemplaris* (Fig. 1B) is established as an emerging model organism in evolutionary developmental biology and extreme tolerance research (5, 6). Like many other tardigrades, it exhibits parthenogenesis and synchronizes its molting period with oviposition to provide additional physical protection to the developing embryos through the exuvium (6, 7). *H. exemplaris* hatch from their eggshells upon completion of embryonic development, measuring about 125 µm in length, and resembling small adults. They progressively increase in size through multiple molts, eventually reaching a fully-grown adult state measuring approximately 240 µm in length ((8), this work). Subsequently, they continue to molt at a relatively constant rate until their death, although these subsequent molting events no longer contribute to additional growth (Fig. 1C). *H. exemplaris* start producing eggs (*i.e.,* reach sexual maturity) shortly after hatching, long before reaching their fully-grown size (Fig.1D; (7, 9)). To simplify the terminology, we named ‘adults’ those tardigrades that had both reached sexual maturity and their fully-grown size, and ‘juveniles’ those that were still growing, irrespective of sexual maturity (Fig. 1C-D).

Tardigrades feature a particular cell type, the storage cells, found floating freely throughout their body cavity fluid ((10, 11); Fig. 1B). Their primary function is to assist in tardigrade nutritional maintenance by storing and distributing energy in the form of protein, glycogen, and fat (11, 12). They also participate in vitellogenesis (11, 13), and potentially contribute to immunity (14). A recent study has shown increased expression of genes belonging to the *SAHS* (secretory abundant heat-soluble) family in storage cells (15), hinting at their potential involvement in desiccation tolerance. In *H. exemplaris*, lipids constitute the primary reservoir material within these cells (11). These lipids can be stained with vital dyes such as BODIPY ((11, 16), see methods), revealing abundant fat in the form of lipid droplets inside the cytoplasm of storage cells (Fig. 1E-F). Storage cells can be isolated from the organism through dissection, enabling independent experimentation with them, separate from the whole organism ((17); Fig. 1F).

Eutely refers to a biological phenomenon observed in certain organisms where somatic cell division ceases once embryonic development is complete, implying a relative constancy in cell numbers throughout an animal’s growth (18). This suggests that subsequent growth involves an increase in cell size rather than an increase in cell count. The prevailing belief, both in peer-reviewed scientific literature (1, 9, 19, 20) and in tardigrade biology outreach websites, is that tardigrades are eutelic animals. However, over 50 years ago, variations in organ cell numbers and potential somatic mitoses among different tardigrade species were already documented (21, 22). More recently, two studies pointed at cell proliferation in two different somatic cell types. One study captured a limited number of storage cells presenting condensed chromosomes using DNA staining approaches in the tardigrade *Richtersius coronifer* (23). Another study showed molecular evidence of cell proliferation at the anterior and posterior ends of the midgut in the tardigrade *H. exemplaris*, a process involved in replacing the cells lining the gut rather than in increasing the overall cell numbers (24). Discrepancies regarding which cell types proliferate were noted by the authors compared to previous claims, possibly due to the starvation conditions in which the experiments were performed (24). In a recent biology book (25), the authors wrote ‘facts as basic as whether or not tardigrades are eutelic required considerable investigation (*i.e.* active bibliographic search) from our side’. Further, in a recent tardigrade review (26) the question ‘Are tardigrades eutelic animals?’ was discussed, and the authors reasoned that this was not the case, due to the isolated indications of cell divisions and the cell number inconstancy studies mentioned above. The ongoing debate surrounding whether tardigrades are genuinely eutelic or not, and the apparent difficulty for both the broader scientific community and the general public to reach a consensus on this matter, underscores the need for a more definitive characterization of the cell types that proliferate under normal physiological conditions within this enigmatic phylum.

In this study, we used *H. exemplaris* to determine whether somatic cell proliferation and an increase in cell numbers occur during tardigrade post-embryonic growth. Our findings reveal that overall cell numbers increase when comparing hatchlings to adults, mainly attributed to a prominent rise in storage cell numbers. Notably, EdU, a molecular tool for monitoring DNA replication, incorporates into storage cells at significantly higher rates during tardigrade growth than in fully-grown adults. We also show that starvation arrests growth and blocks EdU incorporation in storage cells, altogether linking storage cell proliferation to somatic growth. Moreover, we found a direct association between storage cell DNA replication and ecdysis, and reveal that hampering DNA replication provokes animal death during molting. Our results offer a comprehensive insight into DNA replication patterns during tardigrade growth, decisively demonstrating that tardigrades cannot be categorized as eutelic animals.

## RESULTS

### Tardigrade growth is accompanied by an increase in storage cell number

We first collected and fixed animals at both extremes of body size: hatchlings and fully-grown adults (Fig. 1C). We utilized the fluorescent DNA marker DAPI (4’,6-diamidino-2-phenylindole) to quantify the number of cell nuclei present at each growth stage (see methods; Fig. 2A; SI Appendix, Video S1). Our analysis revealed approximately 1140 ± 60 nuclei in hatchlings and approximately 1400 ± 118 in adults, pointing at a significant increase in cell numbers as the animals attained larger sizes (Fig. 2A-B). As we compared adult animals to hatchlings, we observed a notable increase in the area stained by the lipid vital dye BODIPY (Fig. 2C, in green). This led us to hypothesize that this change could be associated with an augmented number of storage cells. Leveraging the nuclei’s position within the animal, their small size, and the intense BODIPY staining surrounding these nuclei, we successfully identified and quantified the number of storage cells in both hatchlings and adult tardigrades. While hatchlings contained 72 ± 10 storage cells, the counts for adult *H. exemplaris* showed they bear 281 ± 64 storage cells, indicating a significant rise in storage cell numbers (Fig. 2C-D; SI Appendix, Video S2). Accordingly, we occasionally observed storage cells that appeared to be in the process of division (SI Appendix, Fig. S1). Intriguingly, the difference in the total number of cells, and the difference in the total number of storage cells between hatchlings and adults was similar, suggesting that the overall increase in cell number primarily results from storage cell proliferation. Our results show that the number of cells increases during tardigrade somatic growth and suggest that storage cells are the main cell type contributing to this raise.

**Figure 2.**
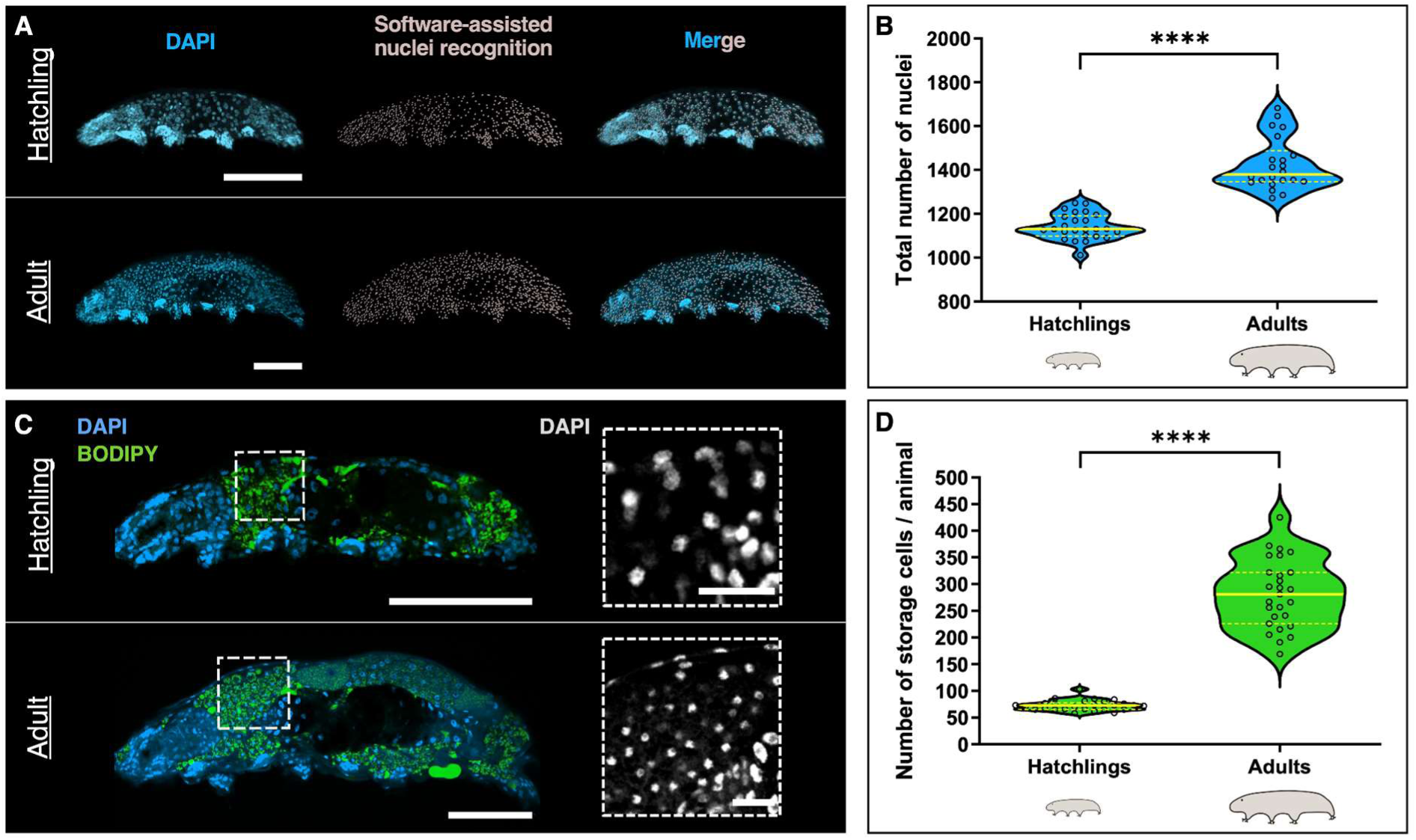
Somatic growth involves overall cell number increase, mainly driven by storage cell number expansion. (A) Representative images of a hatchling (top) and an adult (bottom) of *H. exemplaris*, showing nuclei (DAPI, blue) and software-assisted nuclei recognition (in gray; see methods). Rightmost panels show merged images. The full depth of the animals was captured in confocal z-stacks, and the images shown are maximum projections. (B) Violin plots displaying the total number of nuclei in hatchlings (n = 25) and adults (n = 22). Center yellow lines show the medians; discontinuous yellow lines show the quartiles; the width of the violin at any given point represents the density of data points at that value; each quantified sample is represented by an empty circle. (C) Representative images of a hatchling (top) and an adult (bottom) of *H. exemplaris*, showing DNA (DAPI, blue) and neutral lipids (BODIPY, green). Insets show magnifications of the regions outlined in C, corresponding to the antero-dorsal region of the tardigrade body, were a big proportion of storage cells are normally found (Fig. 1E; (10)). Nuclei in insets are shown in gray. Individual images are shown, rather than maximum projections, to facilitate storage cell visualization within the animals. (D) Violin plots depicting the number of storage cells in hatchlings (n = 26) and adults (n = 27). Violin plot description as in (B). ****, *p* ≤ 0.0001. Scale bars: 50 µm in (A, C); 10 µm in (C insets).

### Storage cells predominantly proliferate during animal growth

To investigate cell proliferation at the molecular level, we monitored DNA replication via the incorporation of the thymidine analog EdU, a marker previously employed successfully in tardigrades (24). Initially, we collected hatchlings and incubated them in EdU for three weeks (Fig. 3A), approximately the duration required for a hatchling to develop into a fully-grown adult when kept at 15°C and fed *ad libitum* (Fig. 1C). This approach had the potential of allowing us to identify any DNA replication that may take place during *H. exemplaris* growth. We detected EdU incorporation in storage cells, germ cells inside the ovary, and gut cells (Fig. 3B-B’). Interestingly, there was an absence of EdU incorporation in any other organ, including epidermal cells, brain cells, ganglia, buccal apparatus, Malpighian tubules, and claw glands (Fig. 3B). While it is known that gut cells incorporate EdU in *H. exemplaris* (24), our observation represents the first molecular evidence demonstrating DNA replication, and thus cell proliferation, in tardigrade storage cells.

**Figure 3.**
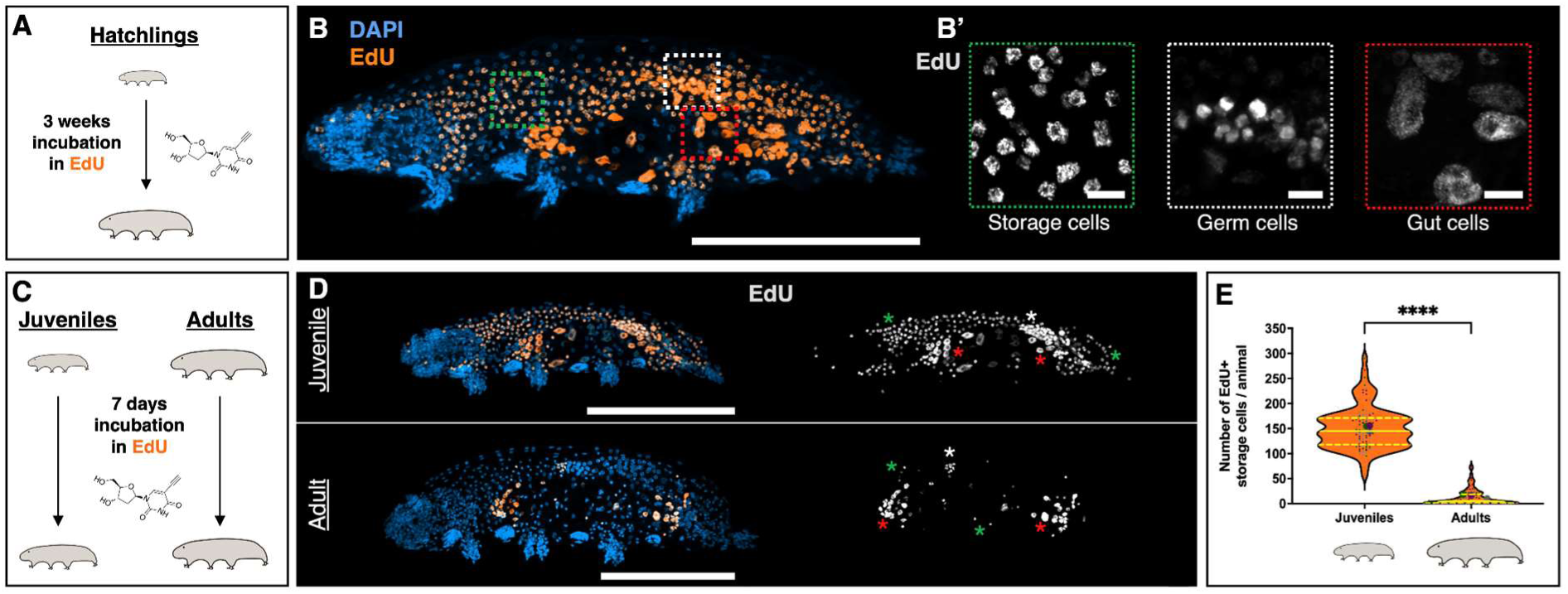
Storage cell proliferation predominantly occurs during somatic growth. (A) Schematic depicting the 3-week EdU incubation experiment, concerning (B). (B) Representative image of tardigrade upon 3-week exposure to EdU showing nuclei (DAPI, blue) and EdU^+^ cells (orange). (B’) Magnification of regions outlined in (B), highlighting the three main replicative cell types labeled with EdU (gray). (C) Experimental schematics for 7-day EdU exposure (D-E). (D) Representative images of juvenile (top) and adult (bottom) after 1-week EdU exposure, displaying nuclei (DAPI, blue) and EdU^+^ cells (orange). Right images show only EdU^+^ cells (gray). (E) Violin plots showing the number of EdU^+^ storage cells in juveniles (n = 95) and adults (n = 90). Plotted values belong to three independent experiments. Each quantified animal is represented by a colored dot, with distinct colors corresponding to individual experiments. Larger dots represent each experiment’s mean. Plot description as in Fig. 2B. Images shown are maximum projections of the animals’ full depth, except for (B’), which focuses on specific cell types using sub-stack projections. Asterisks denote EdU^+^ cell types (storage cells, green; germ cells, white; gut cells, red). ****, *p* ≤ 0.0001. Scale bars: 100 µm in (B, D); 5 µm in (B’).

Tardigrade germ cells reside in the anterior end of the ovary, and derive from primordial germ cells that are specified during embryonic development (27). They form germ cell clusters by undergoing mitosis followed by incomplete cytokinesis (28, 29), thus being expected to incorporate EdU. Within these clusters, the cell completing meiosis develops into an oocyte, while the remaining differentiate into trophocytes (nourishing cells) (28, 29). Accordingly, in addition to EdU incorporation in germ cells, we observed EdU signal in trophocytes and oocytes of *H. exemplaris* (SI Appendix, Fig. S2), corroborating the notion that these cell types are the progeny of germ cell clusters.

Next, we explored whether storage cell proliferation varied between (growing) juveniles and (fully-grown) adults. Juveniles and adults were incubated in EdU for seven days (Fig. 3C), a duration sufficient for all animals to experience at least one molting event when kept at 15°C. We observed that growing juveniles incorporated EdU in a large number of storage cells, whereas we could only detect a limited number of EdU^+^ storage cells in adults (Fig. 3D – green asterisks), rendering the differences highly significant (Fig. 3E). Juveniles’ EdU incorporation in a substantial number of storage cells occurred regardless of the animals’ reproductive status at the time of fixation (non-gravid – Fig. 3D; gravid, or laying eggs – SI Appendix, Fig. S3). Our findings first show that the majority of storage cell number expansion takes place during tardigrade growth. Second, they evidence that storage cell proliferation also happens, although to a much-lessened extent, after tardigrades have reached their fully-grown status.

### Starvation halts animal growth and suppresses storage cell proliferation

Tardigrades exhibit remarkable starvation tolerance, enduring several weeks without food (12). Although prior studies noted that starvation reduces storage cell size (12), little is known about its potential effects on tardigrade growth and cell proliferation. To assess this, we first collected small juveniles of *H. exemplaris* and transferred them to dishes with and without *Chlorococcum* algae (“Fed” and “Starved” conditions). We observed that the majority of starved animals entered a contracted, resting state (Fig. 4A) shortly after food deprivation. Upon reintroducing algae after one week, they resumed activity within minutes, and their guts were filled with algae within 24 hours (SI Appendix, Fig. S4A), indicating reversibility of the ‘starvation state’. We also noticed a marked reduction in lipid staining inside the storage cells of starved individuals after one week (SI Appendix, Fig. S4B), suggesting that the fat reserves of storage cells were utilized to sustain the animals during starvation. Additionally, we collected small juveniles again, measured their length, and exposed them to the aforementioned conditions. After seven days, fed individuals exhibited significant growth, while starved ones showed no substantial increase in size (Fig. 4B-C), implying that starvation induces growth arrest in tardigrades.

**Figure 4.**
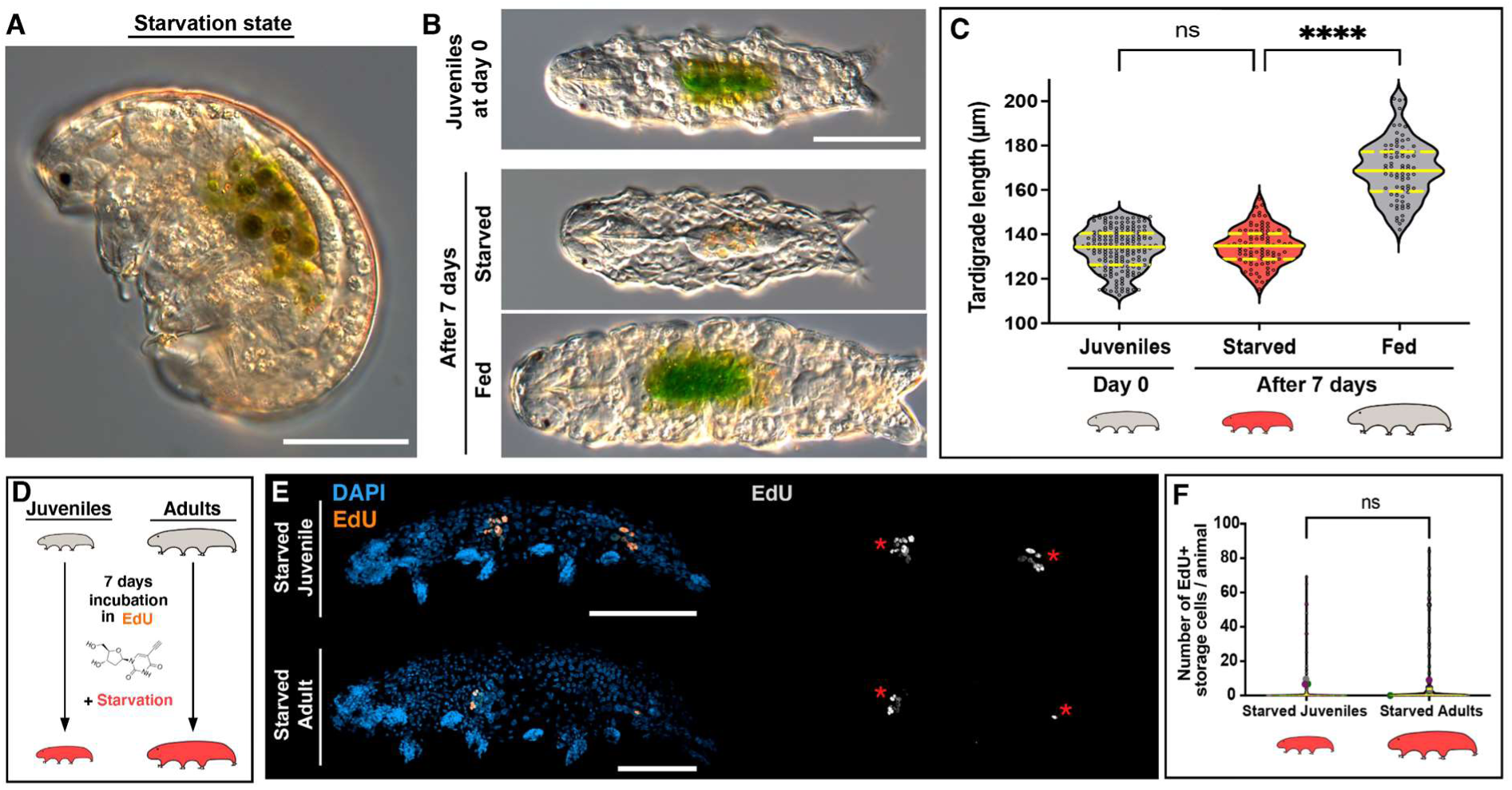
Impact of starvation on growth and storage cell proliferation. (A) Representative image of *H. exemplaris* in the described ‘starvation state’. (B) Representative images of *H. exemplaris* juveniles at day 0 and at day 7 under starved and fed conditions. (C) Violin plots illustrating tardigrade length in juveniles at day 0 (n = 172) and at day 7 under starved (n = 92) and fed (n = 80) conditions. Plot description follows Fig. 2B. (D) Experimental schematics for 7-day EdU exposure in starved juveniles and adults (E-F). (E) Representative images of starved juvenile (top) and starved adult (bottom) after 1-week EdU exposure, showing nuclei (DAPI, blue) and EdU^+^ cells (orange). Right images display only EdU^+^ cells (gray). Red asterisks indicate EdU^+^ gut cells. The full depth of the animals was captured in confocal z-stacks, and the images shown are maximum projections. (F) Violin plots showing the number of EdU^+^ storage cells in starved juveniles (n = 115) and starved adults (n = 112). Plotted values belong to three independent experiments. Plot conventions are consistent with Fig. 3E. Red tardigrade schematics indicate starved condition. ns = non-significant; **** = p ≤ 0.0001. Scale bars: 50 µm.

To assess cell proliferation during food deprivation, we simultaneously exposed juveniles and adults to EdU and starvation for seven days (Fig. 4D). In both cases, we found a complete absence of EdU incorporation in storage cells, with rare exceptions where a few storage cells were EdU^+^ (Fig. 4E-F). Essentially, in most individuals of either growth stage we could only detect EdU^+^ gut cells (Fig. 4E – red asterisks). These results align with a previous study, which showed that starved adult *H. exemplaris* incubated with EdU for up to four days only exhibited EdU incorporation in gut cells (24). Altogether, these findings demonstrate that starvation arrests tardigrade growth and inhibits EdU incorporation in storage cells, unambiguously linking storage cell proliferation to growth.

### EdU incorporation in storage cells takes place during molting

To investigate whether DNA replication in storage cells occurs at specific stages of *H. exemplaris* post-embryonic development or is a continuous process throughout growth, we conducted 24-hour EdU incubations in juvenile animals, including those in proximity to molting (Fig. 5A). The vast majority of animals that did not molt during the 24-hour EdU incubation exhibited no EdU incorporation in storage cells (Fig. 5B-C); only gut cells displayed EdU labeling (red asterisks – Fig. 5B). In contrast, juveniles fixed during molting displayed a significant number of EdU^+^ storage cells (green asterisks in Fig. 5B; Fig. 5C), in addition to EdU^+^ gut and germ cells (red and white asterisks, respectively; Fig. 5B). These results strongly associate storage cell proliferation with the molting process (Fisher’s exact test: *p* < 0.0001; SI Appendix, Table S1). Hence, our findings indicate that storage cell (and germ cell) proliferation does not occur constantly during tardigrade growth but rather in bursts at each ecdysis.

**Figure 5.**
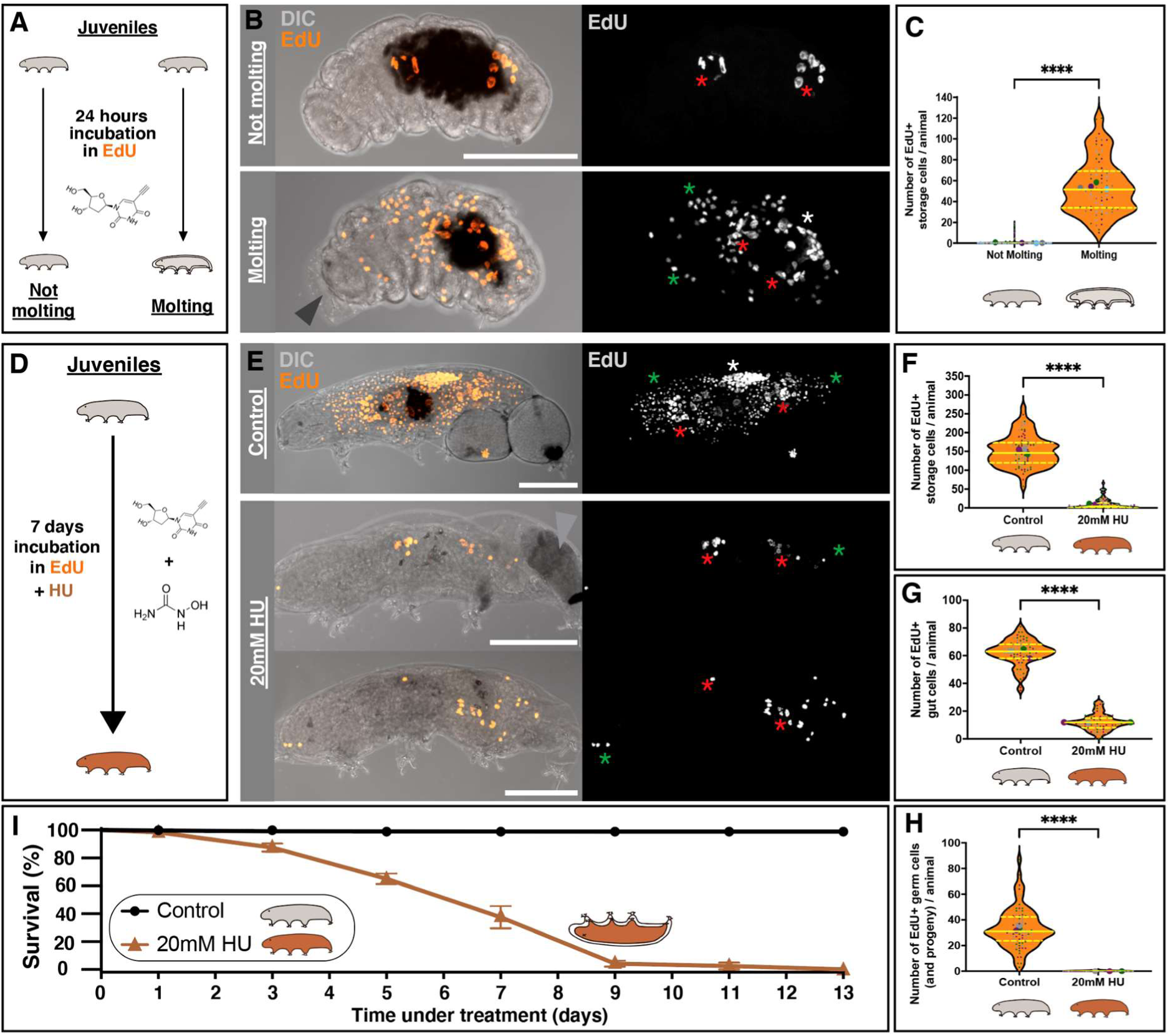
Storage cells undergo replication during molting, while blocking replication results in molting-related fatalities. (A) Experimental schematics for 24-hour EdU exposure (B-C). (B) Representative images of non-molting (top) and molting (bottom) tardigrades upon 24-hour EdU exposure, illustrating animal morphology (DIC, gray) and EdU^+^ cells (orange). Arrowhead indicates the old cuticle being molted. Right images show only EdU^+^ cells (gray). (C) Violin plots displaying the number of EdU^+^ storage cells in non-molting (n = 452) and molting animals (n = 76). Plotted values belong to four independent experiments. (D) Schematic illustrating the 7-day experiment exposing tardigrades to EdU and HU simultaneously, concerning (E-H). (E) Representative images of control (- HU) and 20 mM HU-treated animals after 1-week EdU exposure, showing animal morphology (DIC, gray) and EdU^+^ cells (orange). Arrowhead indicates a laid egg that has degraded. Right images display only EdU^+^ cells (gray). Maximum projections of the animals’ full depth are shown. Asterisks as in Fig. 3. Scale bars = 50 µm. (F-H) Violin plots showing the number of EdU^+^ storage cells (F), gut cells (G), and germ cells (including their progeny: oocytes and trophocytes; H) in control (n = 80) and 20 mM HU-treated animals (n = 93). Plotted values belong to three independent experiments. All plot conventions as in Fig. 3E. ****, *p* ≤ 0.0001. (I) *H. exemplaris* 13-days survival curve in control (-HU) vs 20mM HU conditions (n = 120, across three independent experiments). Means ± standard errors of the mean (SE) are plotted. Brown tardigrade schematics indicate 20mM HU condition. The upside-down brown tardigrade schematic represents animal death during molting.

### Hampering DNA replication progression results in tardigrade death during molting

This prompted us to assess the potential relevance of DNA replication for the molting process. To this purpose, we used hydroxyurea (HU), a drug known to effectively slow down the progression of replication forks and, consequently, EdU incorporation, in other aquatic invertebrates (30, 31). First, to ensure that HU successfully reduces EdU incorporation in replicative cells of *H. exemplaris*, we simultaneously exposed juvenile animals to HU and EdU for seven days (Fig. 5D). We used a 20mM HU dose, intermediate between the doses typically used in yeast (∼100mM) and human cells (1-5mM) (32). We observed an overall decrease in EdU incorporation in animals subjected to 20mM HU compared to control animals (Fig. 5E). Quantification of the number of EdU^+^ storage cells, gut cells, and germ cells (including their progeny: trophocytes and oocytes) in both control and HU-treated condition revealed a highly significant reduction in all cases for animals under HU treatment (Fig. 5F-H). Having confirmed that HU disrupts replication progression in *H. exemplaris* by significantly reducing EdU incorporation in various replicative cell types, we then turned to assess its effects on molting.

In these first experiments, we already observed that, by day 7, a significant number of animals had died. To gain a more comprehensive insight into tardigrade survival under HU exposure, we incubated *H. exemplaris* specimens in 20mM HU and monitored mortality for up to 13 days. After nine days of incubation, allowing sufficient time for all treated animals to molt, we found that 96% ± 4% of the HU-treated animals had perished (Fig. 5I). Remarkably, all dead animals died during ecdysis, irrespective of their size or whether they had undergone egg laying during the process (Fig. 5E), indicating that ecdysis-associated deaths were unrelated to oviposition or egg viability during molting. Altogether, we conclude that hindering DNA replication leads to molting-related fatalities, emphasizing the critical role of accurate DNA replication in allowing effective tardigrade molting and survival.

## DISCUSSION

This study addresses the controversy in the current literature regarding whether tardigrades maintain a constant number of cells during post-embryonic development, and we demonstrate unequivocally that this is not the case. Using the emerging model *H. exemplaris*, we illustrate how tardigrade storage cells primarily undergo proliferation during animal growth, leading to an increase in their numbers. We reveal that food deprivation induces a reversible ‘starvation state’ in *H. exemplaris*, during which both growth and storage cell proliferation are halted. Our 24-hour EdU pulse experiments establish a clear association between molting and storage cell DNA replication, and allowed us to illustrate a comprehensive representation of the DNA replication events occurring during tardigrade post-embryonic life for the three main replicative cell types we identified (Fig. 6). Furthermore, we establish that proper DNA replication is essential for tardigrade survival, as its disruption results in mortality during molting, the sole phase in which gut, germ, and storage cells undergo replication simultaneously (Fig. 6).

**Figure 6.**
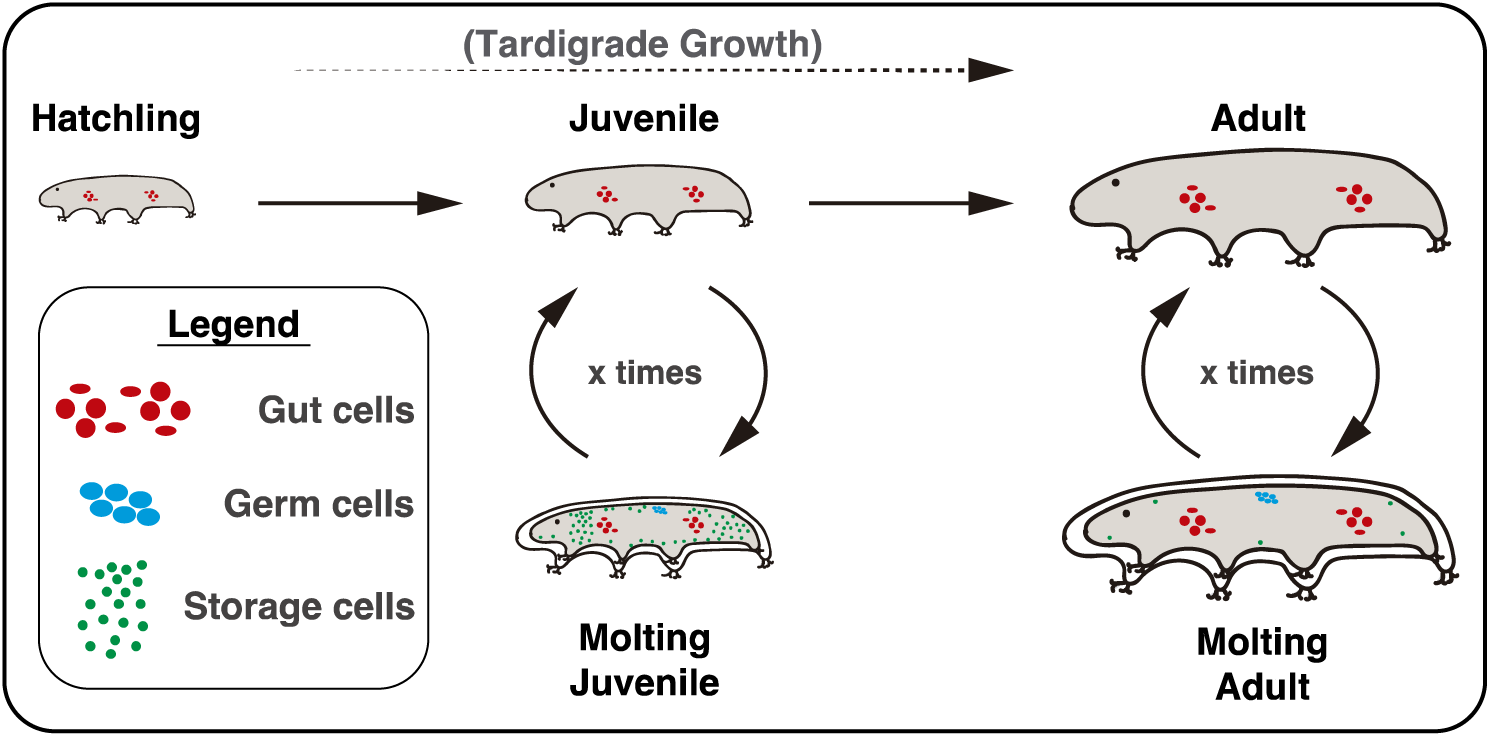
Overview of DNA replication events in tardigrade post-embryonic development. Replicative gut cells are represented as red circles and ovals. Replicative germ cells are represented as blue circles. Replicative storage cells are represented as small green circles. In summary, gut cells in the anterior and posterior parts of the midgut replicate continuously, while germ and storage cells only undergo replication at each molting event. The number of replicative storage cells decreases markedly as tardigrades reach their fully-grown adult size, highlighting that most storage cell proliferation occurs during somatic growth.

*H. exemplaris* does not present a defined number of cells, typical of eutelic animals, as its cell count increases with age and body growth (Fig. 2B). This is reinforced by the observed variability in the total number of cells among individuals of the same species (Fig. 2B). While we have demonstrated that this increase in cell number is primarily attributed to storage cell proliferation (Fig. 2D; Fig. 3), we also detected cell proliferation in the gut and ovary. However, these proliferative events are involved in cell replacement and the formation and sustenance of oocytes, respectively, rather than in overall cell number increase (24, 28, 29). In contrast to observations from decades ago in various species of *Macrobiotus*, where some ganglion and claw gland cells were interpreted as mitotic (21), our investigations did not reveal any evidence of cell proliferation in any other organs or tissues of *H. exemplaris* (Fig. 3B, SI Appendix, Fig. S2). Thus, storage cells represent the primary cell type in *H. exemplaris* that undergoes proliferation and results in a cell number expansion.

Based on our findings, *H. exemplaris* cannot be categorized as entirely eutelic. However, the absence of cell proliferation in specific organs, such as the epidermis and the nervous system, suggests that some degree of cell constancy may exist in various tardigrade organs. To gain a more comprehensive understanding, it would be valuable to reevaluate the proliferative capacities of different cell types in a range of tardigrade species using contemporary molecular techniques like EdU incorporation. This approach could provide insights into whether the absence of cell proliferation in certain organs is a common feature shared across the phylum.

The presence of cell proliferation in a limited number of cell types implies that the growth of most tardigrade organs primarily occurs through an increase in cell size rather than through cell proliferation. In this context, an increment in the number of storage cells may serve as a mechanism to support a larger body size and the associated higher demand for stored materials. This hypothesis has been previously suggested (23), and our data offer two pieces of evidence to support it. First, we show that growth arrest induced by starvation in juvenile tardigrades leads to the inhibition of storage cell proliferation (Fig. 4). Second, in adult *H. exemplaris*, where full growth has been achieved, the majority of storage cells within the animal no longer undergo proliferation (Fig. 3 D-E). From another perspective, considering that storage cells also contribute to vitellogenesis (11, 13), and given that a larger body size allows for a greater number of eggs to develop in the ovary, we postulate that an increase in storage cell numbers throughout tardigrade growth could further enhance the capacity to produce and nourish a larger number of eggs per individual. This aligns with our unreported observations that adult animals consistently produce more eggs than juveniles.

While anterior and posterior midgut cells maintain a constant rate of proliferation (Fig. 6, (24)), germ and storage cells appear to be arrested in G_1_ or in G_o_, only resuming cycling to undergo DNA replication during molting (Fig. 5B-C; Fig. 6). Correspondingly, a similar association between storage cell mitotic activity and molting has been reported in the tardigrade species *R. coronifer* (23). These observations would imply that a specific cue, received at the time of molting, prompts these cell types to resume the cell cycle towards DNA replication. What could be the nature of such a signal? Tardigrades share orthologs of several genes associated with arthropod molting processes, including the ecdysone receptor and early-activated genes driven by ecdysone (33). This indicates a certain level of conservation of molting determinants within the Panarthropoda (33, 34). Notably, recent research has shown that low ecdysone levels can stimulate cell proliferation in *Drosophila* wing imaginal discs (35). Downstream of ecdysteroid hormones, a cascade of evolutionarily ancient ecdysis-related neuropeptides (ERNs) plays crucial roles during ecdysis (34). Originally responsible for regulating life cycle transitions in both molting and non-molting phyla, these ERNs have been co-opted to orchestrate the molting process in ecdysozoans (34, 36). Some of the genes encoding these ERNs are conserved in tardigrades, including eclosion hormone (EH) and crustacean cardioactive neuropeptide (CCAP), which are upregulated at hatching (36). Intriguingly, *H. exemplaris* presents five EH and two CCAP paralogs (36, 37). Along these lines, a neuropeptide-receptor couple has been found to induce meiotic progression in jellyfish oocytes arrested at prophase I (38), suggesting that ERN-receptor couples could also regulate cell cycle progression. Collectively, we propose that ecdysteroid-type molting hormones and/or ERNs may serve as triggers for the resumption of the germ and storage cell cycle in tardigrades.

The question emerges as to why HU kills *H. exemplaris* during molting, irrespective of whether they are juveniles or adults. Indeed, even though adult tardigrades exhibit minimal storage cell proliferation (Fig. 3D-E; Fig. 6), they still perish during molting when exposed to HU treatment. Tardigrade molting events are likely the most energy-demanding stages of their post-embryonic development, since they involve replacing the buccal apparatus, claws, and the entire cuticle (2), synchronization with egg laying (7), and bursts of storage and germ cell proliferation (Fig. 5B-C). Given that gut cells undergo permanent renewal (24), animals’ death during molting could reflect a profound impact of disrupting gut cell proliferation, consequently affecting overall energy absorption. In this scenario, the decrease in gut cell proliferation would have a more deleterious effect on survival than that of storage cells.

Alternatively, yet not exclusively, molting-associated fatalities could be ascribed to more direct effects of HU. HU slows down the progression of replication forks and inhibits the firing of late origins (39). Consequently, the coordination of the replicative program with other DNA-related processes, such as transcription, will be perturbed (40). As shown in other ecdysozoans, ecdysis is a highly transcriptionally-active process (33). Moreover, it has been recently demonstrated that successful molting in the nematode *Caenorhabditis elegans* requires rhythmic accumulation of transcription factors, which, in turn, relies on rhythmic transcription (41). Therefore, HU treatment could enhance tardigrade lethality during molting due to profound alterations in the execution of the ecdysis transcriptional schedule. Additionally, HU induces DNA damage through oxidative stress (42), and also leads to DNA break accumulation when replication fork stalling is long-lasting (43), two DNA damage-associated signals that trigger a robust DNA damage response (44). In the context of *H. exemplaris*, where there is no storage cell proliferation between molting cycles, DNA damage signals may be managed or controlled. However, they may translate into a systemic death signal when this insult occurs during a burst of storage cell proliferation. In agreement, DNA damage in proliferative cell types has recently been shown to trigger an alert in the whole animal that accelerates inflammatory and aging outputs (45).

In summary, our study provides the first molecular evidence of storage cell proliferation in a tardigrade species. It determines that tardigrades do not present cell constancy during post-embryonic development, contrarily to several previous claims, thus resolving a long-standing controversy in the field. Moreover, our data shed new light on tardigrade cell biology and establish a clear connection between tardigrade ecdysis and cell cycle transitions. The finding that storage cells undergo replication opens new possibilities for *in vitro* culture of this cell type and suggests its potential as a valuable model for studying cell cycle dynamics, responses to replication stress, and DNA damage control in tardigrades. With the advent of transgenesis (15) and CRISPR (46) technologies in tardigrades, along with the established RNAi methods (47, 48), *Hypsibius exemplaris* is increasingly becoming a promising genetic experimental model, altogether marking an exciting era in tardigrade research.

## MATERIAL AND METHODS

### Animal husbandry and drug treatment

*H. exemplaris* (Z151 strain) husbandry was conducted as previously described (49), with minor modifications. In brief, tardigrades were maintained in an incubator at 15 °C, placed inside 55 mm diameter plastic Petri dishes filled halfway with spring water (Volvic) filtered through a 0.2 µm mesh. To facilitate tardigrade movement, the bottom of the dishes was scratched with sandpaper. Photoperiod and relative humidity inside the incubator were not monitored. Animals were fed *ad libitum* with *Chlorococcum* algae, cultivated in 50 ml tubes with BG-11 Growth Media (Gibco, A1379901). Water changes were performed every two weeks. All experimental incubations lasting ≥24 hours were carried out under these same conditions in 35 mm diameter dishes, placed inside cardboard boxes to avoid degradation of potentially photosensitive compounds.

Tardigrades were subjected to incubation with the ribonucleotide reductase inhibitor hydroxyurea (HU; Sigma-Aldrich, H8627) at a concentration of 20 mM, dissolved in filtered spring water (FSW) for the duration of the experiment. Renewal of HU took place every 2-3 days to ensure its continued effectiveness.

### DAPI and BODIPY staining

Animals were collected and filtered through a 40 µm mesh, then transferred to a glass depression slide within a humid chamber using a glass Pasteur pipette. Whenever possible, samples were kept in darkness for the duration of the experiment. To label neutral lipids, live specimens were incubated in 10 µg/ml BODIPY (Difluoro{2-[1-(3,5-dimethyl-2H-pyrrol-2-ylidene-N)ethyl]-3,5-dimethyl-1H-pyrrolato-N}boron; Sigma-Aldrich, 790389) diluted in FSW for 30 minutes. Subsequently, samples were rinsed three times with FSW and fixed in 4% PFA in 1x PBS for 1 hour at room temperature (RT). After fixation, three 15-minute washes in 1x PBS were performed. Samples were then incubated in the DNA fluorescent stain DAPI (4’,6-diamidino-2-phenylindole; Sigma-Aldrich, D9542) at a concentration of 1 µg/ml in 1x PBS for 30 minutes. Following this step, the samples were washed three times in 1x PBS for 15 minutes, transferred to a slide, and mounted in ProLong Gold Antifade Mountant (Invitrogen, P36930) prior to confocal imaging.

### EdU experiments

To detect DNA replication, animals were incubated in 100 µM EdU (EdU Click-iT™ Cell Proliferation Kit for Imaging, Alexa Fluor 488 Dye; Invitrogen, C10337) diluted in FSW containing algae (except during experimental starvation) for specific durations (24 hours, 7 days, or 3 weeks). In the 3-week experiment, EdU was refreshed weekly. Following EdU exposure, tardigrades were collected, transferred to a glass depression slide, and fixed with 4% PFA in 1x PBS + 1% Triton X-100 (PTx) for 1 hour at RT. After fixation, the samples underwent three 15-minute washes in PTx. Subsequently, they were blocked for 2 hours in a 0.2 µm-filtered blocking solution containing 10% Bovine Serum Albumin (BSA) in 1x PBS. The Click-iT EdU detection reaction was carried out for 1 hour at RT according to the manufacturer’s instructions. After the detection reaction, four 10-minute PTx washes were conducted. DAPI was used to stain nuclei, and samples were mounted in ProLong Gold for confocal imaging, as described above. In some cases, tardigrades were euthanized with 10% EtOH before fixation to prevent the animals from contracting during exposure to PFA.

### Imaging, length measurements, cell counting and statistics

To capture live images of tardigrades, animals were anesthetized using 20mM levamisole hydrochloride (MedChemExpress, HY-13666) in mqH_2_O and mounted on slides, covered with coverslips. Clay was positioned in each corner of the coverslip to prevent compression, except for the intentionally compressed specimen in Fig. 1B. Differential interference contrast (DIC) color images were taken with a Leica K3C digital color camera attached to a compound light microscope (Leica Thunder). For measuring the body length of tardigrades, measurements were conducted using ImageJ (50). The measurement was taken from the head to the juncture on the posterior-most segment with legs (9). For starved tardigrades, measurements were obtained immediately after animals came back to an active state, ensuring their full body extension.

Fluorescently labeled images were acquired using a Zeiss LSM980 confocal microscope, equipped with an Airyscan2 module. To ensure consistency when comparing samples within a given experiment, identical scanning parameters were applied to all conditions in each independent experiment. Confocal Z-stacks and projections were processed and adjusted for brightness and contrast in ImageJ, using K. Terretaz’s visualization toolset (https://github.com/kwolbachia/Visualization_toolset).

To count the total number of cells in hatchlings and adults, Imaris software (Oxford instrument) was employed. Nuclei were highlighted using custom thresholding to allow for quantification. Quantifications of BODIPY-stained storage cells and EdU^+^ cells were performed manually in ImageJ. In all instances, z-stacks encompassing the entire depth of the animals were utilized. Supplementary videos S1 and S2 were assembled using Imaris and ImageJ software, respectively. Tardigrade schematics and figure compilation were generated using Adobe Illustrator.

To determine statistical significance, normality of all datasets was first assessed using the Shapiro-Wilk test. Two-tailed Student’s *t*-test was conducted for the dataset concerning total number of storage cells (Fig. 2D). For all other comparisons, Mann-Whitney *U* nonparametric tests were employed for two-way comparisons, and the Kruskal-Wallis test was used for statistical analyses in Fig. 4C. Fisher’s exact test was selected for the association assessment, based on a 2 × 2 contingency table. Statistical significance for all quantitative comparisons is represented as ****, where *p*-value ≤ 0.0001. Graphs and statistical analyses were performed using GraphPad Prism 9.

## Supporting information

Video S1 - Software-assisted nuclei recognition in H. exemplaris

Video S2 - Storage cell number increase in adult animals

## Abbreviations

DAPI: (4’,6-diamidino-2-phenylindole),
BODIPY: (Difluoro{2-[1-(3,5-dimethyl-2H-pyrrol-2-ylidene-N)ethyl]-3,5-dimethyl-1H-pyrrolato-N}boron),
EdU: (5-ethynyl-2’-deoxyuridine),
HU: (hydroxyurea).

## ACKNOWLEDGEMENTS

We thank Simon Galas and Myriam Richaud for providing us with *H. exemplaris* and *Chlorococcum* samples, and for giving us advice on tardigrade manipulation. We also thank Benjamin Lacroix for helping us to establish the model by sharing his equipment, and Kseniya Samardak and Sylvain Kumanski for their help with tardigrade husbandry. We are grateful to Kevin Terretaz for his valuable tips and macros on ImageJ. We also express gratitude to the CRBM direction for its support. We acknowledge the imaging facility MRI, member of the France-BioImaging national infrastructure supported by the French National Research Agency (ANR-10-INBS-04, «Investments for the future»). This research was supported by the I-SITE MUSE (Montpellier Université d’Excellence).

## AUTHOR CONTRIBUTIONS

Conceptualization: G.Q.A. and M.M.C

Formal Analysis: G.Q.A.

Funding acquisition: M.M.C

Investigation: G.Q.A.

Methodology: G.Q.A.

Project administration: G.Q.A. and M.M.C

Supervision: M.M.C

Validation: G.Q.A.

Visualization: G.Q.A.

Writing – original draft: G.Q.A. and M.M.C

Writing – review & editing: G.Q.A. and M.M.C

The authors declare no competing interest.

## Notes

### Competing Interest Statement

The authors have declared no competing interest.

