## Supplementary Information for "Storage cell proliferation during somatic growth establishes that tardigrades are not eutelic organisms"

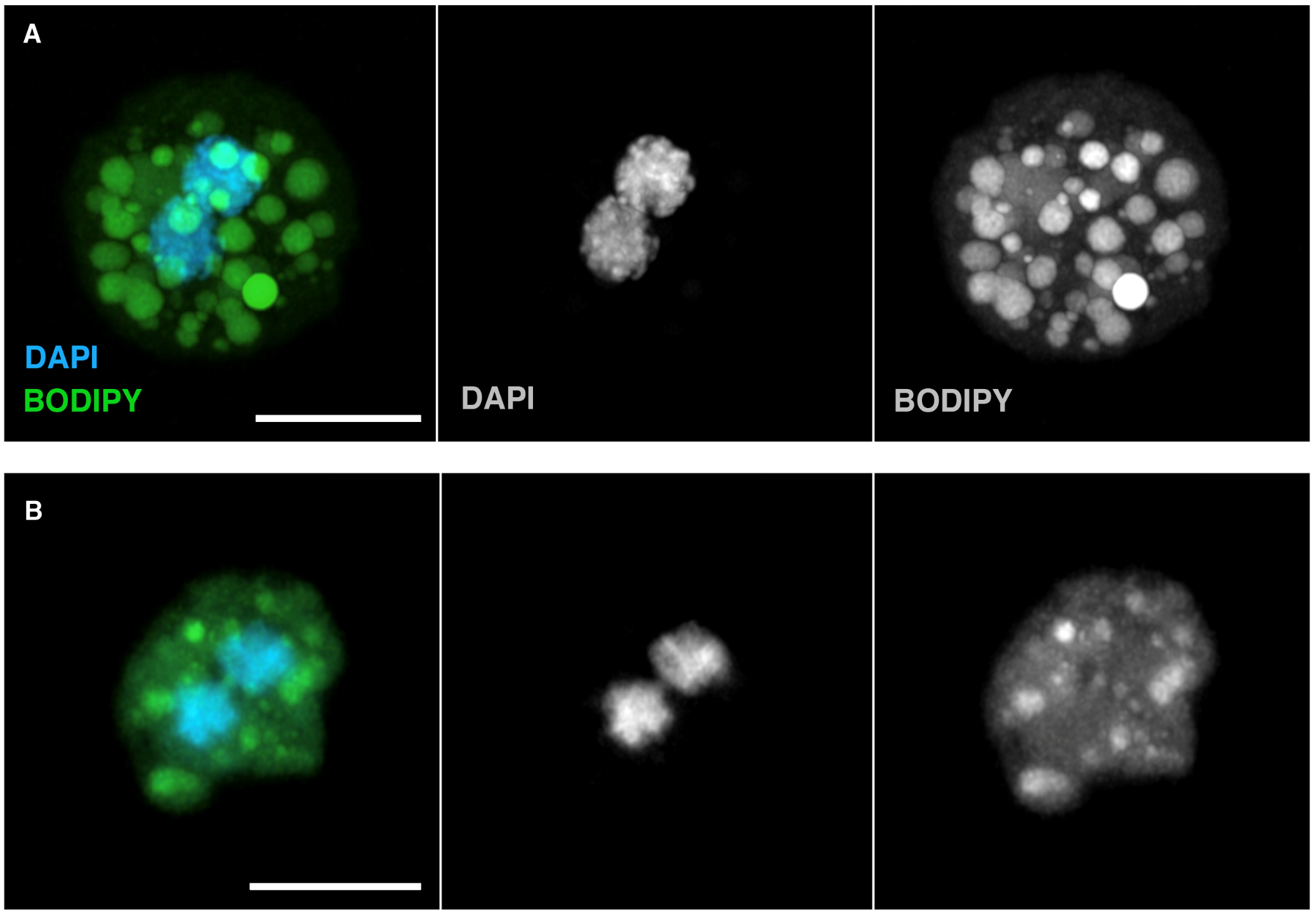

**Supplementary Figure S1. Potentially dividing storage cells.** (A-B) Two instances of storage cells, originating from different juvenile *H. exemplaris* specimens, imaged outside the animal. Both examples exhibit the presence of two nuclei within the storage cells, suggesting these cells were captured during / after mitosis and / or before cytokinesis. Nuclei (DAPI) are depicted in blue, and neutral lipids (BODIPY) are shown in green. Images shown are maximum projections of the storage cells’ full depth. Scale bars = 5 µm.

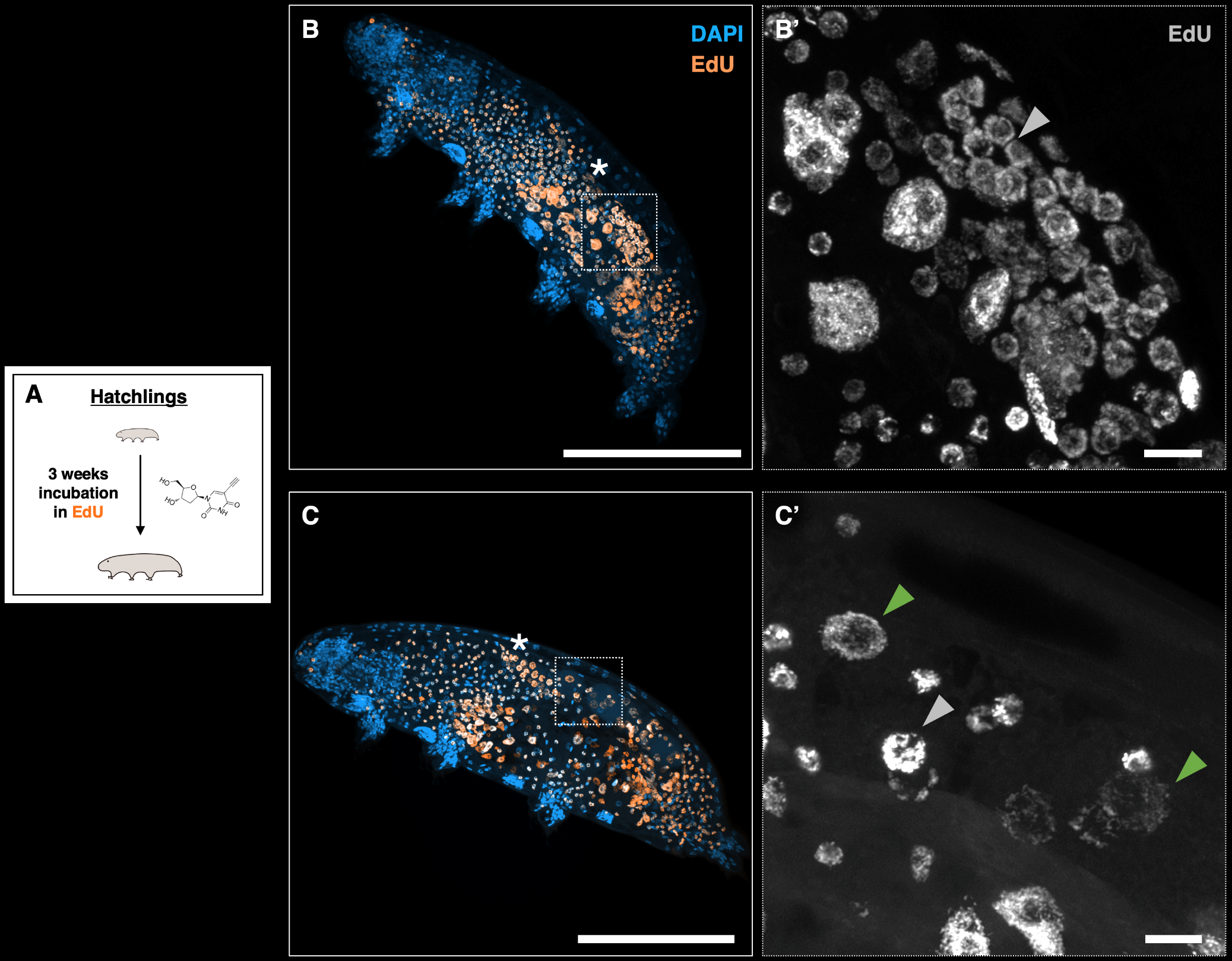

**Supplementary Figure S2. EdU^+^ trophocytes and oocytes.** (A) Schematic illustrating the 3-week EdU incubation experiment relevant to (B-C). (B-C) Two different images of *H. exemplaris* after a 3-week exposure to EdU, depicting nuclei (DAPI, blue) and EdU^+^ cells (orange). (B) represents a tardigrade in the early oogenesis stage, while (C) shows a gravid tardigrade containing developing oocytes. White asterisks in (B, C) indicate EdU^+^ germ cells. (B’) Magnification of the region outlined in (B), focusing on an area of the ovary full of EdU^+^ trophocytes. (C’) Magnification of the region outlined in (C), highlighting an area of the ovary with EdU^+^ oocytes. In (B’-C’), EdU is displayed in gray. Green arrowheads indicate EdU^+^ oocyte nuclei, while gray arrowheads point to EdU^+^ trophocyte nuclei. The images in (B, C) are maximum projections of the animals’ full depth, while those in (B', C') are sub-stack projections to enhance the visualization of the specific cell types of interest. Scale bars: 100 µm in (B, C); 5 µm in (B’, C’).

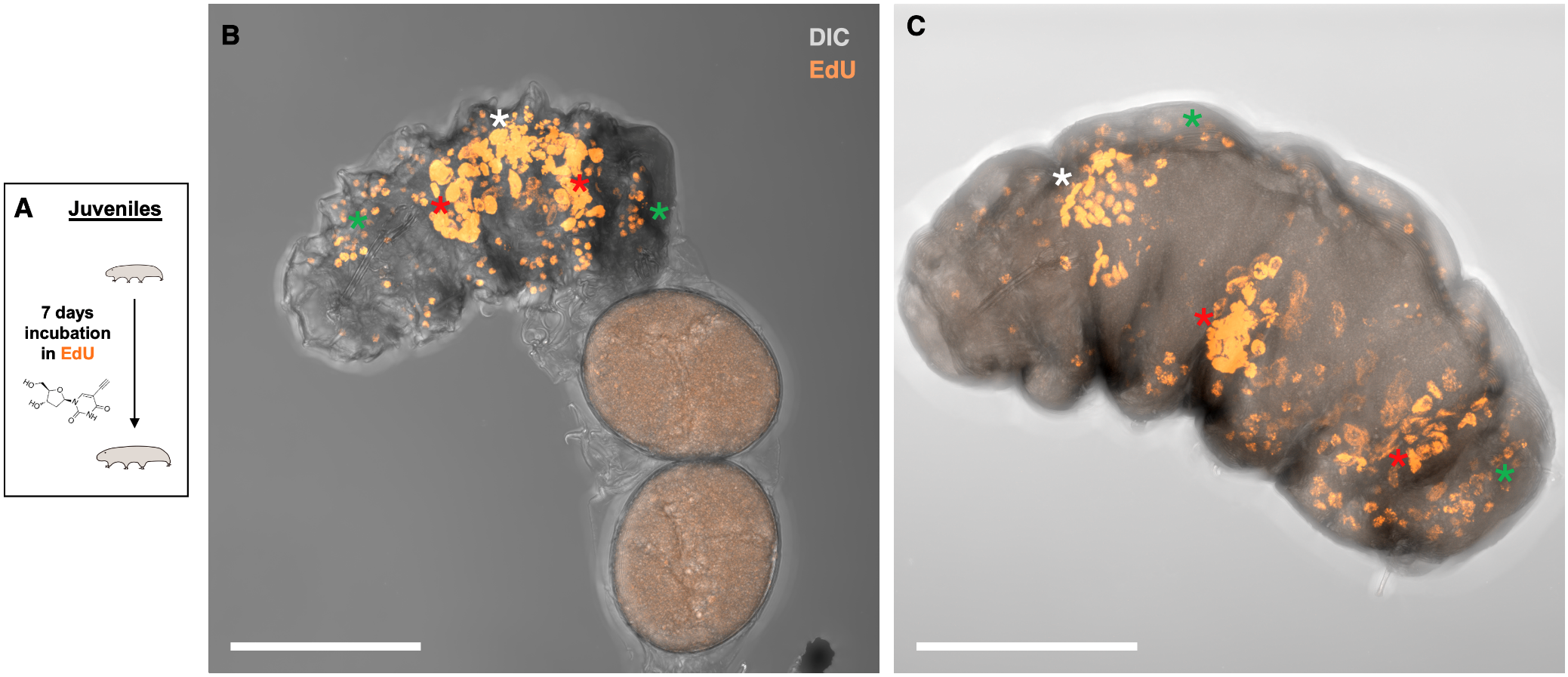

**Supplementary Figure S3. Storage cell proliferation in egg-laying and gravid juveniles.** (A) Schematic illustrating the 7-day experiment exposing tardigrades to EdU relevant to (B-C). (B) Image of a juvenile tardigrade fixed after egg laying, and about to leave the exuvium behind, following 1-week exposure to EdU. (C) Image of gravid juvenile after 1-week exposure to EdU. Animal morphology (DIC) is depicted in gray, and EdU^+^ cells are shown in orange. Maximum projections of the animals’ full depth are presented. Asterisks as indicated in Fig. 3. Scale bars = 50 µm.

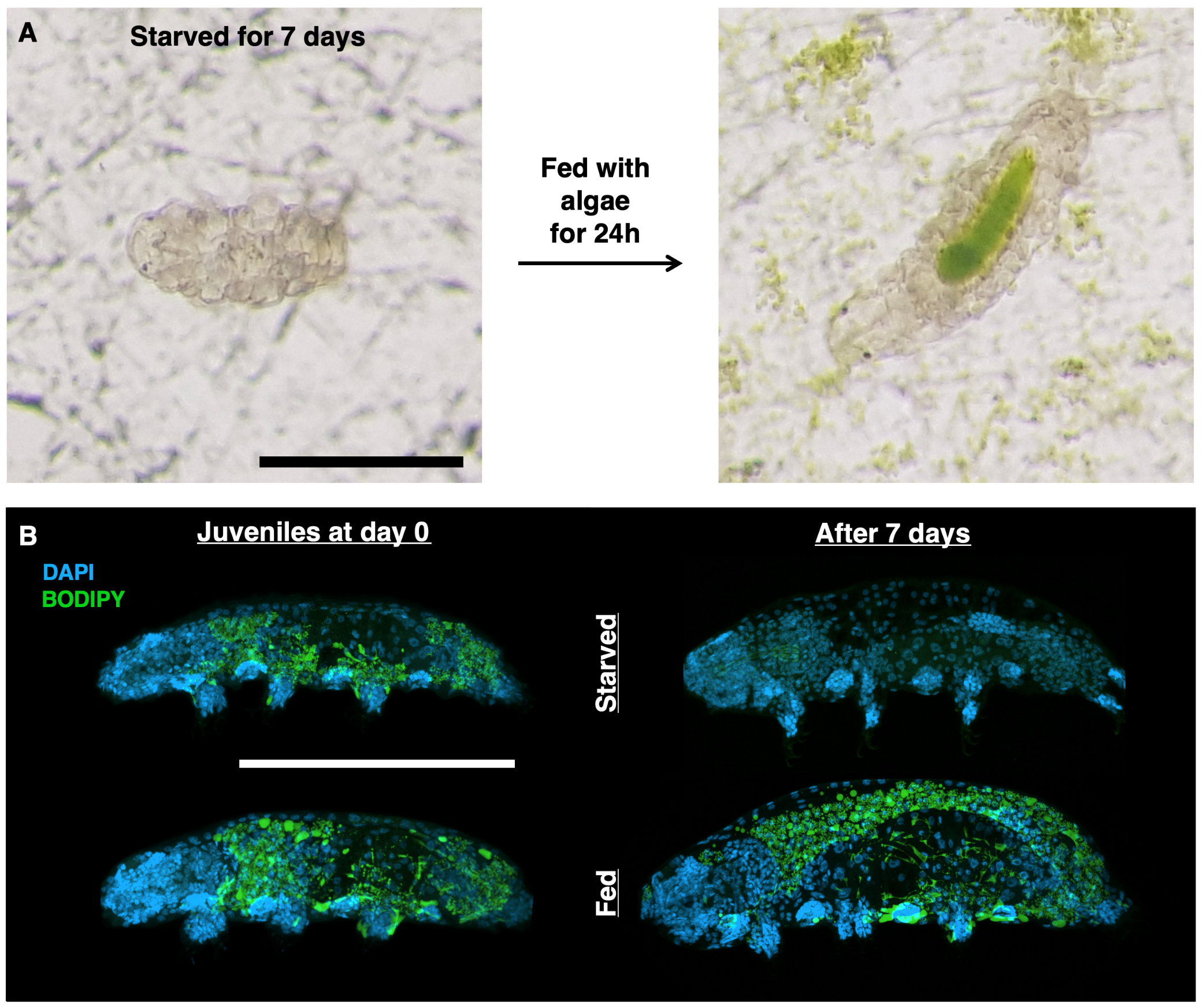

**Supplementary Figure S4. Reversibility of tardigrade ‘starvation state’ and loss of storage cells’ lipids after 1-week starvation.** (A) Images of a tardigrade that has been starved for one week (left) and upon addition of *Chlorococcum* algae for 24 hours (right). ­(B) Representative images of juveniles at day 0 and at day 7 under starved and fed conditions, showing DNA (DAPI, blue) and neutral lipids (BODIPY, green). Note the nearly complete absence of BODIPY staining in starved animals after 7 days of starvation. The full depth of the animals was captured in confocal z-stacks, and the images shown are maximum projections. Scale bars: 100 µm.

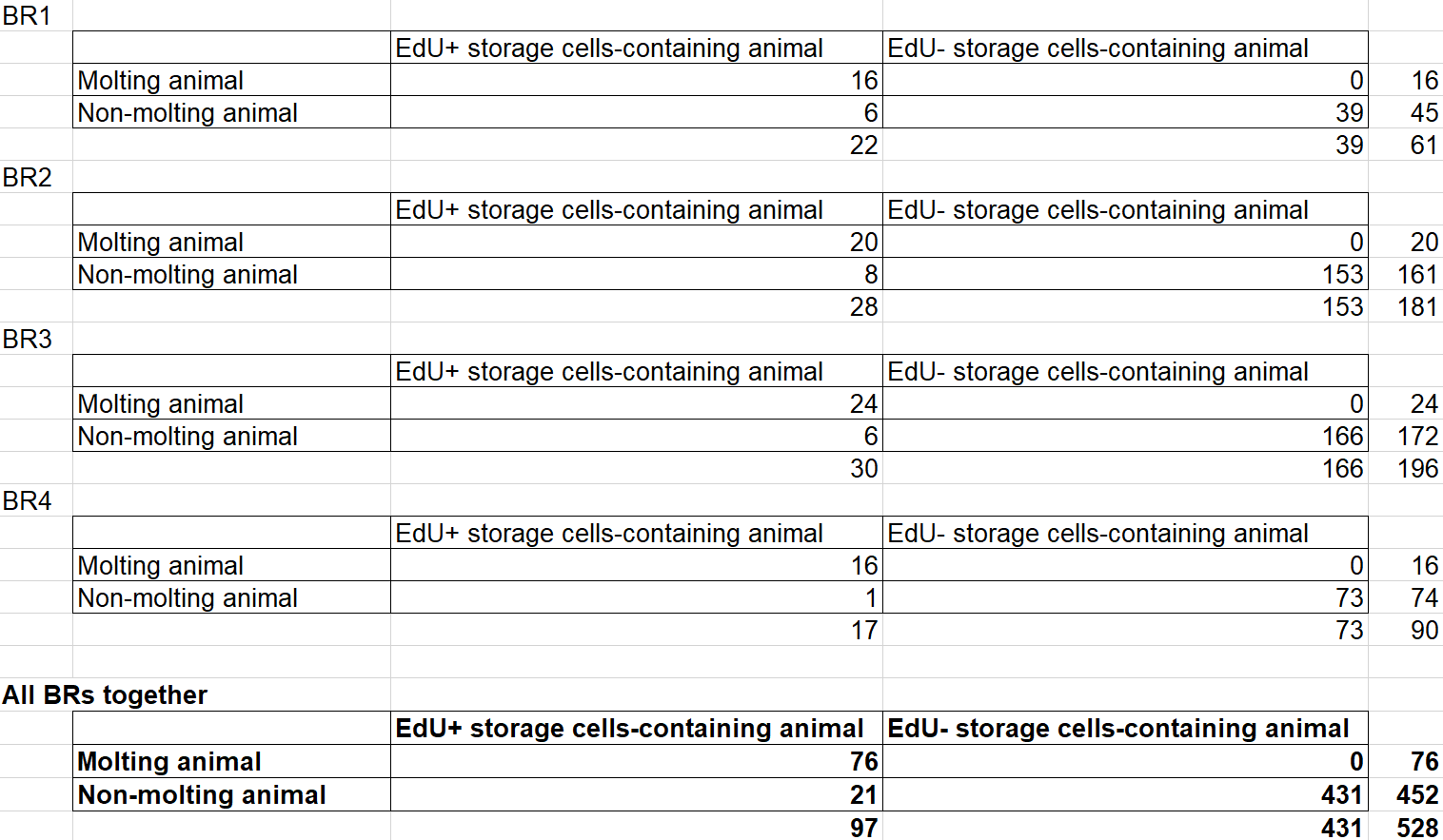

**Supplementary Table S1. Association of storage cell proliferation with ecdysis.** This table summarizes data from four independent experiments examining the association between the presence of EdU^+^ storage cells and whether animals were molting or not. Note: EdU^+^ storage cells-containing animals were counted as positive as soon as at least one storage cell was found to be EdU^+^. The two-tailed Fisher's exact test was performed on the 2 × 2 contingency table pooling the data from the four biological replicates (BR), resulting in a p-value lower than 0.0001. Therefore, the association between rows and columns is considered extremely significant.

**Supplementary Video S1. Software-assisted nuclei recognition in *H. exemplaris*.** Representative hatchling and adult specimens processed in Imaris software for nuclei recognition and number quantification. Nuclei are shown in blue, and software-assisted nuclei recognition is displayed in gray.

**Supplementary Video S2. Storage cell number increase in adult animals.** This video showcases the same hatchling and adult specimens of *H. exemplaris* as shown in Fig. 2C. It presents a z-stack of each animal image by image, with a 1 µm interval between pictures. The video is presented at the same scale to emphasize the size differences and the significant increase in storage cell numbers when comparing adult and hatchling tardigrades. Nuclei are displayed in blue, and neutral lipids are shown in green.
